# Pattern separation in cerebellar-targeted whisking pre-motor nuclei

**DOI:** 10.1101/2025.09.16.676487

**Authors:** Staf Bauer, Peipei Zhai, Nathalie van Wingerden, Vincenzo Romano

## Abstract

The cerebellar output can trigger whisker movement through indirect projections that pass via several brainstem pre-motor nuclei before reaching the facial nucleus, which directly controls whisker movements in rodents. Although the central pattern generator function of the intermediate reticular formation has been recently clarified, the roles of the other whisker pre-motor nuclei remain unclear. Here, we set out to compare the whisker movement kinematics of the main pre-motor whisker nuclei connecting the cerebellum and the facial nucleus. We optogenetically stimulated neurons located in the cerebellar cortex Purkinje cells (PCs), the cerebellar nuclei (CN), the red nucleus (RN), the superior colliculus (SC), the spinal trigeminal nucleus (SV), and the reticular formation (RF); in head-fixed awake mice while monitoring the bilateral whisker movement. Here, we find that optogenetic stimulation of the RN, SC, and SV resulted in a predominant midpoint change, whereas optogenetic stimulation of the PCs, CN, and RF resulted in faster whisker movements. In addition, the excitation of PCs, the RN, and SC resulted in symmetric bilateral whisking. In contrast, the excitation of CN, the RF, and SV resulted in initial asymmetric movement, followed by a more dominantly symmetrical bilateral whisking. Our results suggest that several pre-motor nuclei receiving cerebellar output modulate different aspects of whisking, suggesting pattern separation takes place in the brain-wide network downstream of the cerebellum.

## 1. Introduction

Different animal species utilize diverse forms of active sensing to explore their environment. Rodents actively protract and retract their whiskers to collect sensory information from their surroundings (Kleinfeld, Ahissar, & Diamond, 2006; Prescott, Diamond, & Wing, 2011). The whisker system in rodents has been used as a standard model to unravel sensorimotor control (Auffret et al., 2018; Crochet, Poulet, Kremer, & Petersen, 2011; Hill, Curtis, Moore, & Kleinfeld, 2011; Lang, Sugihara, & Llinas, 2006; Lindeman et al., 2021; O’Connor, Berg, & Kleinfeld, 2002; Romano et al., 2022; Zhai et al., 2024). Rodents have specialized muscles to control whisker movements (Bellavance et al., 2017; Freeman & Dale, 2013; Hill, Bermejo, Zeigler, & Kleinfeld, 2008). While intrinsic muscles directly affect the fast component of the whisker movements at each cycle, that of the extrinsic muscles largely determines their slow component of the midpoint (Bellavance et al., 2017; Hill et al., 2008). These muscles are innervated by different groups of motor neurons located in the facial nucleus (FN), which receives input from pre-motor nuclei like the red nucleus (RN), reticular formation (RF), the superior colliculus (SC),, and the spinal trigeminal nucleus (SV) (Bosman et al., 2011; Hattox, Priest, & Keller, 2002).

The RN, SC, RF, and SV are reciprocally connected with the cerebellum (Bosman et al., 2011; May, 2006; Novello, Bosman, & De Zeeuw, 2022; Takatoh et al., 2022; T.M. Teune, J. van der Burg, J. van der Moer, J. Voogd, & T. J. Ruigrok, 2000). Optogenetic excitation of Purkinje cells has been shown to induce bilateral whisking with different forms of symmetry (Romano et al., 2022; Zhai et al., 2024). Research groups investigating the neural circuit controlling rhythmic whisker behaviour have made big advancements in unravelling the role of the RF as a central pattern generator for whisking (Deschenes, Takatoh, et al., 2016; Golomb et al., 2022; Kleinfeld, Deschenes, Wang, & Moore, 2014; Moore et al., 2013; Sreenivasan, Karmakar, Rijli, & Petersen, 2015; Takatoh et al., 2022). However, whisker behaviour is state-dependent and occurs quasi-rhythmically with varying amplitude, midpoint, and frequency. In addition, whisker behaviour can consist of non-periodic movements that precede head and body movements (Dominiak et al., 2019; Sofroniew, Cohen, Lee, & Svoboda, 2014; Towal & Hartmann, 2006). Therefore, whether other pre-motor nuclei, which are reciprocally connected to the cerebellum and project to the FN, contribute to different aspects of whisking, and to which, is still unclear. To determine the contribution of the brain areas connecting the cerebellum and FN on whisking kinematics, we optogenetically stimulated PCs, CN, RN, SC, RF, and SV neurons and simultaneously tracked the bilateral whisker movement in head-fixed, awake mice. We show here that stimulation of pre-motor nuclei induces heterogeneous bilateral whisker movements. Excitation of PCs in the medial Paramedian lobule (PML) results in prolonged rhythmic whisking, while stimulation of excitatory and inhibitory cerebellar nuclei (CN) can results in a heterogeneous bilateral whisker pattern. Additionally, stimulation of the RF and SV induces bilaterally asymmetrical whisker movements while stimulation of the SC and RN induces bilaterally symmetrical movements. Finally, we compared the frequencies at which mice whisked during optogenetic cerebellar stimulation versus those of pre-motor nuclei. The power spectrum related to whisking during cerebellar stimulation and pre-motor nuclei revealed that they both cover the same range of frequencies, which are similar to the natural range of whisking frequencies. Beyond the frequency domain, a temporal pattern separation was evident in the time-frequency domain. These results highlight the role of several pre-motor nuclei involved in the brain-wide network downstream of the cerebellum that controls the whisker muscles. The dissimilar spatiotemporal whisking features encoded by the different pre-motor nuclei suggest that pattern separation takes place downstream of the cerebellum.

## 2. Methods

### 2.1 Mice

All experiments were done according to the Animal guidelines of the institutional animal welfare committee of Erasmus MC by the Central Authority for Scientific Procedures. Wild-type C57BL/6J (No. 000664), transgenic Ai27D (No. 012567), and transgenic Ai39 (No. 014539) mice were obtained from the Jackson Laboratory. Mice of 6-34 weeks old were used in this study and mice were housed individually in a 12-hour light-dark cycle with food and water ad libitum. The ambient housing temperature was maintained at ∼25.5 °C with 40-60% humidity. We used 55 mice for the optogenetic stimulation experiments and sacrificed 16 of them for anatomical experiments and histological examination of transgene expressions.

To express ChrimsonR specifically in excitatory or inhibitory CN neurons, AAV9-Syn-FLEX-ChrimsonR-tdTomato was injected in the CN of VGluT2-Cre or Gad2-Cre mice, respectively.

### 2.2 Virus injections and optic fibre placement

AAV9-Syn-ChrimsonR-tdTomato were obtained from UNC Vector Core. All viral vectors were aliquoted and stored at -80**°**C until used. To express ChrimsonR in CN, RN, SC, RF, and SV, 80nl of AAV9-Syn-ChrimsonR-tdTomato viral vectors were injected in the right CN (A-P: -2.7 mm, M-L: 0.8 mm D-V: -2.3 mm relative to lambda) and SV (A-P: -1.55 mm, M-L: 1.8 mm D-V: -3.75 mm relative to lambda), left RN (A-P: -3.5 mm, M-L: 0.9 mm D-V: -3.5 mm relative to bregma), SC (A-P: -3.6 mm, M-L: 1.8 mm D-V: -2.0 mm relative to bregma), and RF (A-P: -3.0 mm, M-L: -0.9 mm D-V: -3.5 mm relative to lambda). After 6 weeks of incubation, an optic fibre was implanted (200µm in diameter, Thorlabs, Newton, NJ, USA) for direct stimulation of the corresponding virus injection region (CN, RN, SC, RF, and SV, respectively) approximately 1 mm above the injection coordinate. After a survival period of ∼10 weeks, the animals were sacrificed and their brains were processed for histology.

### 2.3 Surgeries

For all mice, a magnetic pedestal was placed on the skull above the bregma using Optibond adhesive (Kerr Corporation, Orange, CA) as described before (Romano et al., 2022). A craniotomy was made over the right paramedian lobule (PML) of the cerebellum in 6 mice used for optogenetic stimulation of PCs. Isoflurane anaesthesia (Pharmachemie, Haarlem, The Netherlands; 2-4% V/V in O_2_) was maintained during the whole surgery procedure. Mice were given 5 mg/kg carprofen (“Rimadyl”, Pfizer, New York, NY), 50 µg/kg buprenorphine (“Temgesic”, Reckitt Benckiser Pharmaceuticals, Slough, United Kingdom), 1 µg lidocaine (AstraZeneca, Zoetermeer, The Netherlands), and 1 µg bupivacaine (Actavis, Parsippany-Troy Hills, NJ, USA) to reduce post-surgical pain. After three days of recovery, mice were habituated to the recording apparatus for about 45 minutes during at least 2 daily sessions. During stimulation experiments, mice were head-fixed with the pedestal and restrained.

For the stimulation of the CN, RN, SC (3 out of 7), RF, and SV, a 2-4 mm long optical fiber (Φ = 200 µm; 0.22 NA, Thorlabs) was implanted through a small cranial window (Φ = 300 µm) and chronically fixed to the skull with dental cement. The optical fibre was placed approximately 1 mm above the viral injection site (see 2.2 for coordinates).

### 2.4 Whisker movement recording and tracking

Whisker movement was recorded and tracked as previously described in (Bauer et al., 2022; Romano et al., 2022; Zhai et al., 2024). A high-speed camera (acA640-750 um, Basler Electric, Highland, IL, USA) was placed ∼50 cm above the mouse to videorecord the whiskers. Frames were captured with 750 Hz and the whisker movements were tracked using our new software tracking tool (Betting et al., 2020), (https://gitlab.com/c7859/neurocomputing-lab/whisker/whiskeras-2.0) and resampled to 1000 Hz using the ‘resample’ function in MATLAB R2022b. Whisker movements were described as the average angle of all trackable whiskers per frame, as previously described in (Bauer et al., 2022; Romano et al., 2022).

### 2.5 Optogenetic stimulation and control experiments

For PC stimulation, LED photo-excitation (wavelength = 595 nm, M595F2, Thorlabs, Newton, NJ, USA) or photo-inhibition (wavelength = 474 nm, M474F2, Thorlabs, Newton, NJ, USA) was given by a high-power light driver (DC2100, Thorlabs, Newton, NJ, USA) through an optical fibre (400 µm in diameter, Thorlabs, Newton, NJ, USA). The optical fibre was placed on the surface of the right PML. For CN, RN, SC (3 out of 7), RF, and SV stimulation, the optical fibre (200 µm in diameter, Thorlabs, Newton, NJ, USA) was connected to the implanted fibre inside the brain. LED photo-excitation was given by the same high-power light driver (DC2100, Thorlabs, Newton, NJ, USA). To be consistent with previous studies, our stimulation lasted 100ms with 1-2s intervals, similar to (Proville et al., 2014; Zhai et al., 2024). The intensities varied between 1 and 1000 mA. The intensity was calibrated to trigger whisker movements with the lowest possible intensity. When stimulation yielded no response, the intensity was increased up to a maximum of 1000 mA (resulting in ∼10 mW). We have previously demonstrated that the impact of such stimulation in Cre-negative mice, that did not express the channel rhodopsin, does not cause any whisker movement (Romano et al., 2020). To exclude also the possibility that LED stimulation picked up by the retina caused whisker movements, we performed control experiments by placing an external optical fibre (400 µm in diameter, Thorlabs, Newton, NJ, USA) in front of the mouse, using the same intensity (Figure S2). During all our control experiments, except those for inhibitory CN stimulation, we see that the control stimulation yields no response, as can be seen in the flat average whisking traces (Figure S2). We used varying intensities between 1 and 1000 mA, calibrated to determine to trigger responses. Because stimulation of inhibitory CN did not trigger whisker movements consistently, during these experiments we used the highest intensity (i.e., 1000 mA). Such strong light stimulation elicited some late whisker responses which were different from those elicited by stimulation of the pre-motor nuclear neurons (Figure S2).

### 2.6 Histology and microscopy

As described previously (Romano et al., 2022), animals were deeply anaesthetized with isoflurane and injected with pentobarbital sodium solution (50 mg/kg) intraperitoneally. Transcardial perfusion was performed with saline, followed by 4% paraformaldehyde (PFA) in 0.1 M phosphate buffer (PB, pH 7.4). Brains were removed immediately and post-fixed for an hour in 4% PFA in 0.1 M PB. Fixed brains were placed in 10% sucrose overnight at 4**°**C and then embedded in 12% gelatin-10% sucrose. After fixation in 10% formalin for an hour, the blocks were placed in 30% sucrose overnight at 4**°**C, 40 µm serial coronal sections were cut with a freezing microtome (SM2000R, Leica) and collected in 0.1 M PB. For immunofluorescence, sections were incubated subsequently with primary and secondary antibodies. All antibodies were titrated for working solution in a 2% normal horse serum-0.4% triton-0.1 M PBS solution. Tissue was incubated in primary antibodies at 4**°**C overnight and in secondary antibodies at room temperature for 2 h. After each incubation with antibodies, sections were gently rinsed with 0.1 M PBS (10 min, 3 times) and subsequently mounted for microscopy. For all immunofluorescence sections, DAPI was used for general background staining. For fluorescence imaging, we took overviews of the brains with a 10x objective on a fluorescence scanner (Axio Imager.M2, ZEISS).

### 2.7 Statistics and visualization

For Figure 1, each mouse received stimulations on the medial PML (excitatory stimulation: 7 mice, inhibitory stimulation: 6 mice), for Figure 2 to Figure 6, each mouse received direct stimulations on CN (Figure 2, excitatory stimulation: 7 mice, inhibitory stimulation: 8 mice), RN (Figure 3, 9 mice), SC (Figure 4, 7 mice), RF (Figure 5, 4 mice), and SV (Figure 6, 7 mice), respectively. The baseline of the induced whisker movement was calculated by the average movement of the same location from all the mice from -200 ms to -100 ms before the stimulation. Peak protraction or retraction angles that exceeded baseline plus or minus 3 times the standard deviation, respectively, were considered movements. Throughout the manuscript, the whisker heatmap was plotted using a custom code based on the MATLAB function “imagesc”. We chose this function because it allows for the simultaneous visualisation of three variables in a 2D plot. Specifically, we used it to show the change in whisker angle at each trial at each time point. This is particularly useful to show how consistent the whisker movement is across trials. The average whisker movements from the example mouse are the average of all the trials, and the SEM is calculated among all the trials. The average whisker movements of one stimulation situation are calculated among all the mice that got stimulation of the same site.

**Figure 1:**
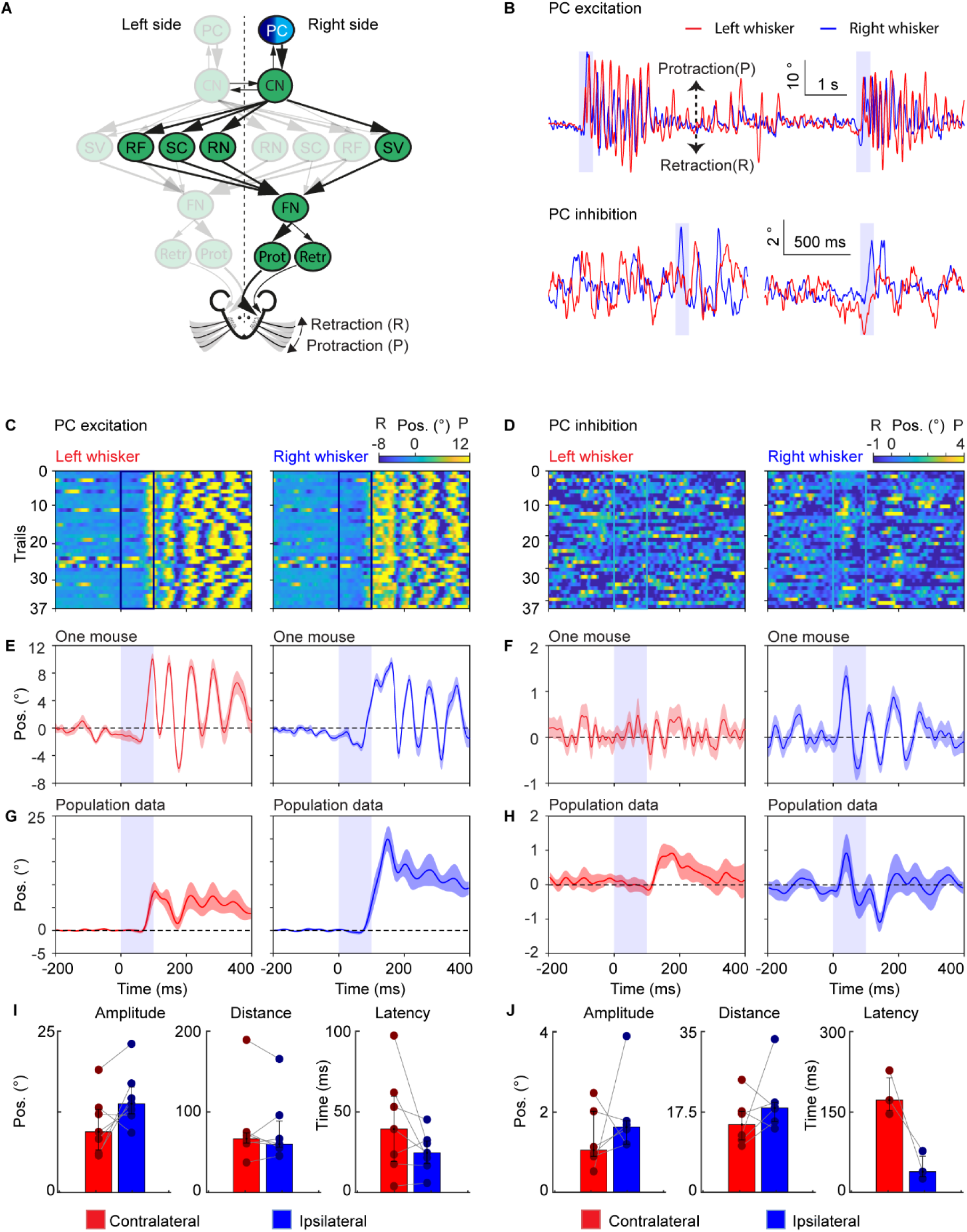
The activation of Purkinje cells, rather than their inhibition, triggers a highly rhythmic pattern of bilateral whisker movement. (**A**) A schematic of the neural circuit downstream of the cerebellar cortex to the facial nucleus. Dark and light blue indicates excitation and inhibition of PCs. (**B**) Two traces of representative profiles of whisker movements upon excitation (top) and inhibition (bottom) of PCs in the medial Paramedian lobule (PML). (**C**) Whisker movements around excitation of PCs in the medial PML in one mouse over 37 trials. Whisker movements around the stimulation of each trial are shown as heat maps. Positive (yellow) and negative (blue) values indicate protraction and retraction, respectively. The colored frames (dark blue in C and bright blue in **D**) represent the optogenetic stimulation pulse of 100 ms; each row shows the average of all whiskers in a single trial. (**D**) Similar to (**C**) during optogenetic inhibition of PCs. (**E**) The average left and right whisker movements of the mouse in (**C**). (**F**) Similar to (**E**), the average whisker trace of the mouse is shown in (**D**). (**G**) The average whisker movements of 7 mice during optogenetic excitation. (**H**) The average whisker movements of 6 mice during inhibition of PCs. The purple shaded column represents the 100 ms optogenetic stimulation. The shaded areas around the lines represent the SEM. (I) Bar plots showing the amplitude, distance and latency after the onset of stimulation during PC excitation. (**J**) Similar to (**I**) during PC inhibition. Both amplitude and distance show the value in degrees and the latency is shown in milliseconds. Bar plots represent median +/- interquartile ranges and individual datapoints. Abbreviation: Purkinje cell of the Paramedian lobule (PC), cerebellar nuclei (CN), facial nucleus (FN), reticular formation (RF), superior colliculus (SC), spinal trigeminal nucleus (SV), the red nucleus (RN), protractor muscles (Prot), retractor muscle (Retr), angle whisker position (Pos).

**Figure 2:**
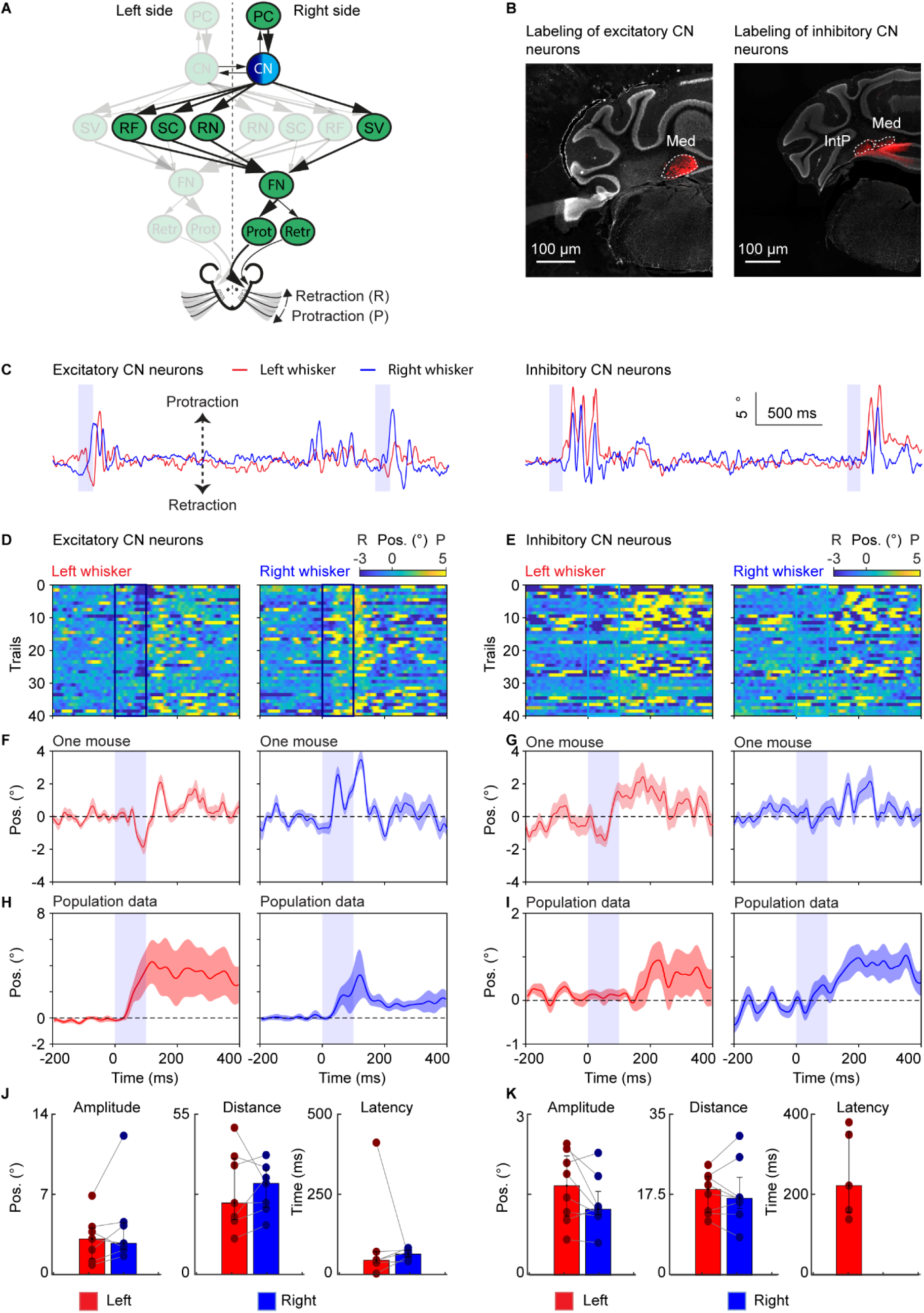
Neither stimulation of excitatory nor inhibitory cerebellar nuclear neurons alone trigger a rhythmic pattern of bilateral whisker movement. (**A**) A schematic of the neural circuit downstream of the cerebellar cortex to the facial nucleus. Dark and light blue indicates stimulation of excitatory and inhibitory CNs, respectively. (**B**) Histological examination of transgene expression in excitatory (left) and inhibitory (right) CN neurons. Med: medial cerebellar nucleus, IntP: interposed cerebellar nucleus. (**C**) Two traces of representative profiles of whisker movements upon stimulation of excitatory (left) and inhibitory (right) CN neurons. (**D**) Whisker movements around stimulation of excitatory CN neurons over 40 trials in one mouse. Whisker movements for each trial are shown as heatmap plots in this example mouse. One row indicates one trial. Positive (yellow) and negative (blue) values indicate protraction and retraction, respectively. The dark blue square represents the period of stimulation. (**E**) Similar to (**D**) but for stimulation of inhibitory CN neurons. Light blue indicates the period of stimulation. (**F**) The average contralateral (left) whisker movements of the mouse in (**D**) respectively. (**G**) Average right whisker movement, similar to (**F**). (**H**) and (**I**) are similar to (**F**) and (**G**), but it’s the average whisker movements of 7 mice during stimulation of excitatory CN neurons (**H**) and 8 mice during stimulation of inhibitory (**I**) CN neurons. The purple shaded column represents the 100 ms duration of optogenetic stimulation. The shaded areas around the lines represent the SEM. (**J**) Bar plots showing the amplitude, distance and latency after the onset of stimulation during PC excitation. (**K**) Similar to (**J**) during PC inhibition. Both amplitude and distance show the value in degrees and the latency is shown in milliseconds. Bar plots represent median +/- interquartile ranges and individual datapoints. Abbreviation: Purkinje cell of the Paramedian lobule (PC), cerebellar nuclei (CN), facial nucleus (FN), reticular formation (RF), superior colliculus (SC), spinal trigeminal nucleus (SV), the red nucleus (RN), protractor muscles (Prot), retractor muscle (Retr), angle whisker position (Pos).

**Figure 3:**
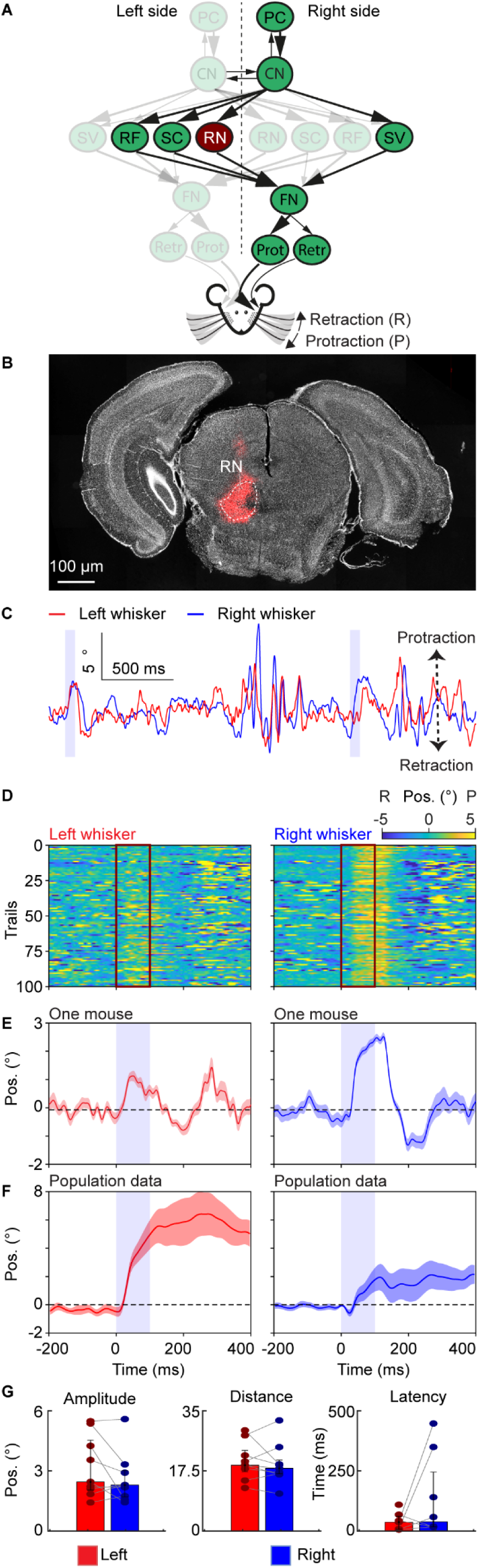
Neuronal excitation in the red nucleus triggers bilaterally symmetrical non-rhythmic whisker movement. (**A**) A schematic of the neural circuit downstream of the cerebellar cortex to the facial nucleus. Dark red indicates the site of stimulation, the RN. (**B**) Histological examination of transgene expression in the RN. RMC: red nucleus, magnocellular part. (**C**) One trace of representative profiles of whisker movements upon stimulation of the RN. The purple shaded column represents the 100 ms optogenetic stimulation. (**D**) Whisker movements of the left and right side around excitation of RN cells over 100 trials in one mouse. Whisker movements around the stimulation of each trial are shown as heatmap raster plots. Positive and negative values indicate protraction and retraction, respectively. The dark red frames represent the period of stimulation and each row shows whisker movement in a single trial. (**E**) The average left and right whisker movements of the mouse in (**D**). (**F**) is similar to (**E**), but it’s the average whisker movements of 9 mice. The shaded areas around the lines represent the SEM. (**G**) Bar plots showing the amplitude, distance and latency after the onset of stimulation during excitation of the RN. Bar plots represent median +/- interquartile ranges and individual datapoints. Abbreviation: Purkinje cell of the Paramedian lobule (PC), cerebellar nuclei (CN), facial nucleus (FN), reticular formation (RF), superior colliculus (SC), spinal trigeminal nucleus (SV), the red nucleus (RN), protractor muscles (Prot), retractor muscle (Retr), angle whisker position (Pos).

**Figure 4:**
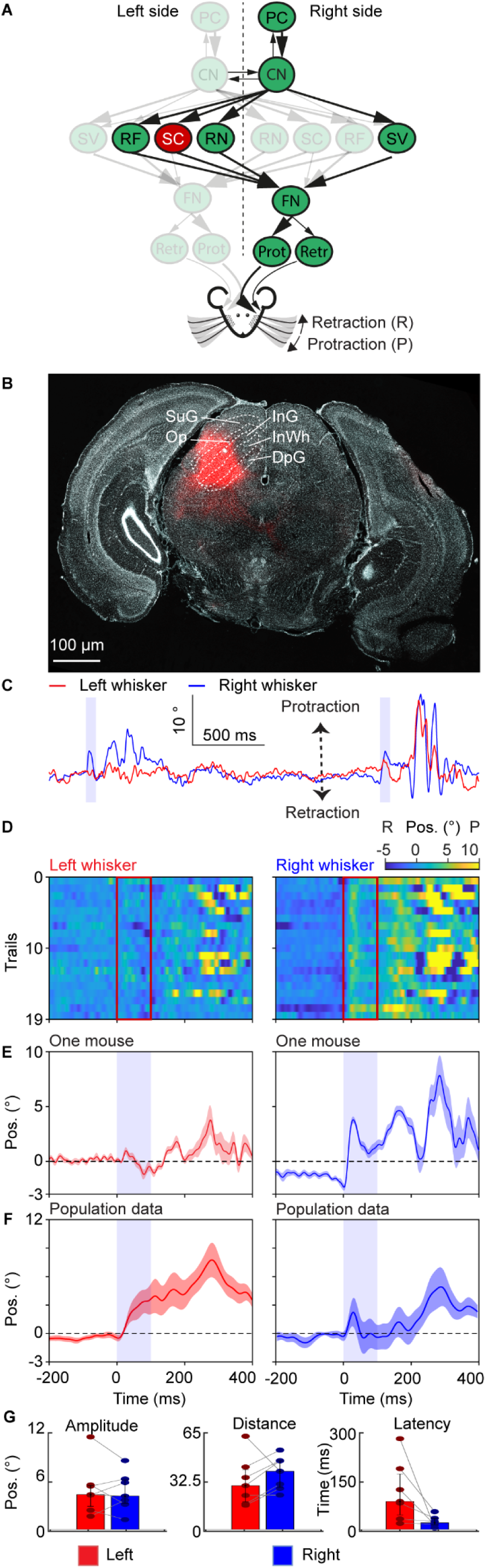
Neuronal excitation in the superior colliculus triggers bilaterally symmetrical quasi-rhythmic whisker movement. (**A**) A schematic of the neural circuit downstream of the cerebellar cortex to the facial nucleus. Red indicates the site of stimulation, the SC. (**B**) Histological examination of transgene expression in the SC. InG: the intermediate grey layer of the superior colliculus, InWh: the intermediate white layer of the superior colliculus. DpG: the deep grey layer of the superior colliculus, Op: optic verve layer of the superior colliculus, SuG: the superficial grey layer of the superior colliculus. (**C**) One trace of representative profiles of whisker movements upon stimulation of the SC neurons. The purple shaded column represents the 100 ms optogenetic stimulation. (**D**) Whisker movements of the left and right side around stimulation of SC cells over 19 trials in one mouse. Whisker movements around the stimulation of each trial are shown as heatmap raster plots. Positive and negative values indicate protraction and retraction, respectively. The red frames represent the duration of stimulation, each row shows whisker movement in a single trial. (**E**) The average left and right whisker movements of the mouse in (**D**). (**F**) is similar to (**E**), but it’s the average whisker movements of 7 mice. The shaded areas around the lines represent the SEM. (**G**) Bar plots showing the amplitude, distance and latency after the onset of stimulation during excitation of the SC. Bar plots represent median +/- interquartile ranges and individual datapoints. Abbreviation: Purkinje cell of the Paramedian lobule (PC), cerebellar nuclei (CN), facial nucleus (FN), reticular formation (RF), superior colliculus (SC), spinal trigeminal nucleus (SV), the red nucleus (RN), protractor muscles (Prot), retractor muscle (Retr), angle whisker position (Pos).

**Figure 5:**
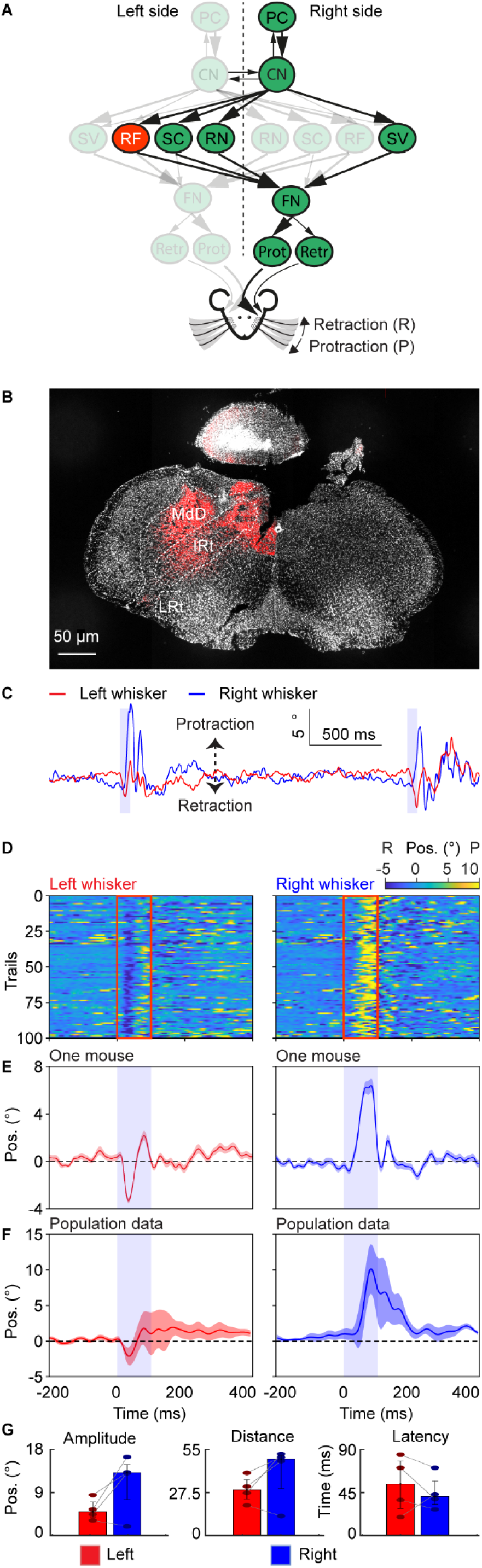
Neuronal excitation in the reticular formation triggers highly asymmetrical fast bilateral whisker movement. (**A**) A schematic of the neural circuit downstream of the cerebellar cortex to the facial nucleus. Red indicates the site of stimulation, the RF. (**B**) Histological examination of transgene expression in the intermediate reticular nucleus (IRt). (**C**) One trace of representative profiles of whisker movements upon stimulation of IRt. The purple shaded column represents the 100 ms optogenetic stimulation. (**D**) Whisker movements of the left and right side around stimulation of IRt cells over 100 trials in one mouse. Whisker movements around the stimulation of each trial are shown as heatmap raster plots. Positive and negative values indicate protraction and retraction, respectively. The red frames represent the stimulation duration, each row shows whisker movement in a single trial. (**E**) The average left and right whisker movements of the mouse in (**D**). (**F**) is similar to (**E**), but it’s the average whisker movements of 4 mice. The shaded areas around the lines represent the SEM. (**G**) Bar plots showing the amplitude, distance and latency after the onset of stimulation during excitation of the RF. Bar plots represent median +/- interquartile ranges and individual datapoints. Abbreviation: Purkinje cell of the Paramedian lobule (PC), cerebellar nuclei (CN), facial nucleus (FN), reticular formation (RF), superior colliculus (SC), spinal trigeminal nucleus (SV), the red nucleus (RN), protractor muscles (Prot), retractor muscle (Retr), angle whisker position (Pos).

**Figure 6:**
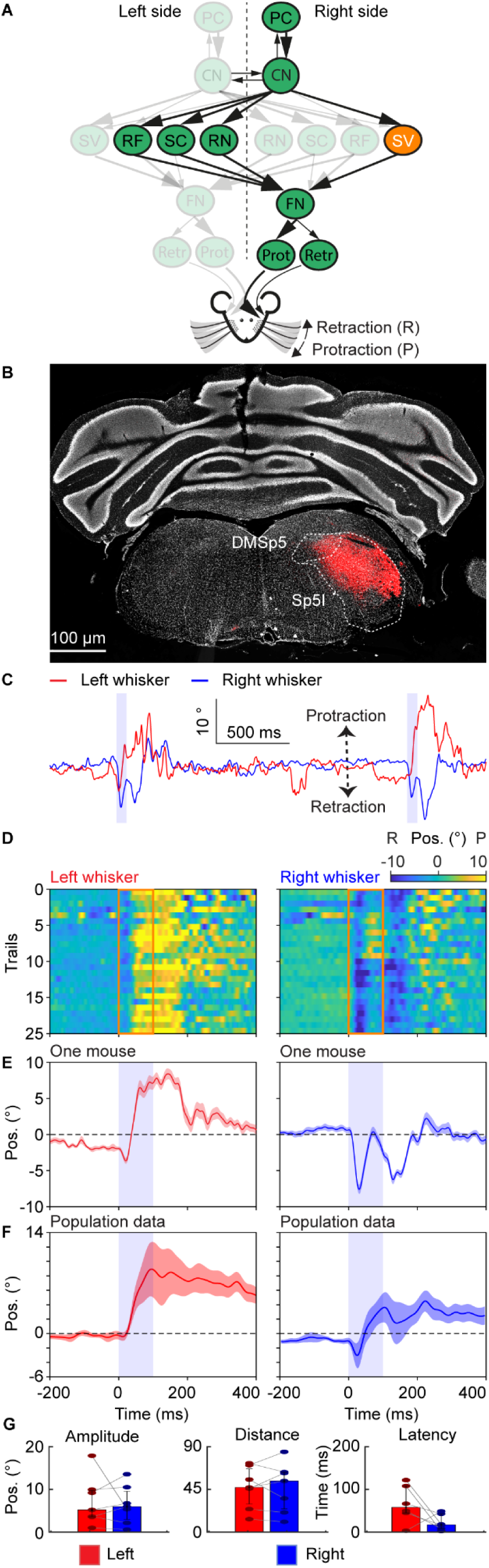
Neuronal excitation in the spinal trigeminal triggers highly asymmetrical slow bilateral whisker movement. (**A**) A schematic of the neural circuit downstream of the cerebellar cortex to the facial nucleus. Orange indicates the site of stimulation, the SV. (**B**) Histological examination of transgene expression in SV. DMSp5: dorsomedial spinal trigeminal nucleus, Sp5I: spinal trigeminal nucleus, interpolar part. (**C**) One trace of representative profiles of whisker movements upon stimulation of SV. The purple shaded column represents the 100 ms optogenetic stimulation. (**D**) Whisker movements of the right and left side around stimulation of SC cells over 25 trials in one mouse. Whisker movements around the stimulation of each trial are shown as heatmap raster plots. Positive and negative values indicate protraction and retraction, respectively. The orange frames represent the stimulation duration, each row shows whisker movement in a single trial. (**E**) The average right and left whisker movements of the mouse in (**D**). (**F**) is similar to (**E**), but it’s the average whisker movements of 7 mice. The shaded areas around the lines represent the SEM. (**G**) Bar plots showing the amplitude, distance and latency after the onset of stimulation during excitation of the SV. Bar plots represent median +/- interquartile ranges and individual datapoints. Abbreviation: Purkinje cell of the Paramedian lobule (PC), cerebellar nuclei (CN), facial nucleus (FN), reticular formation (RF), superior colliculus (SC), spinal trigeminal nucleus (SV), the red nucleus (RN), protractor muscles (Prot), retractor muscle (Retr), angle whisker position (Pos).

**Figure 7:**
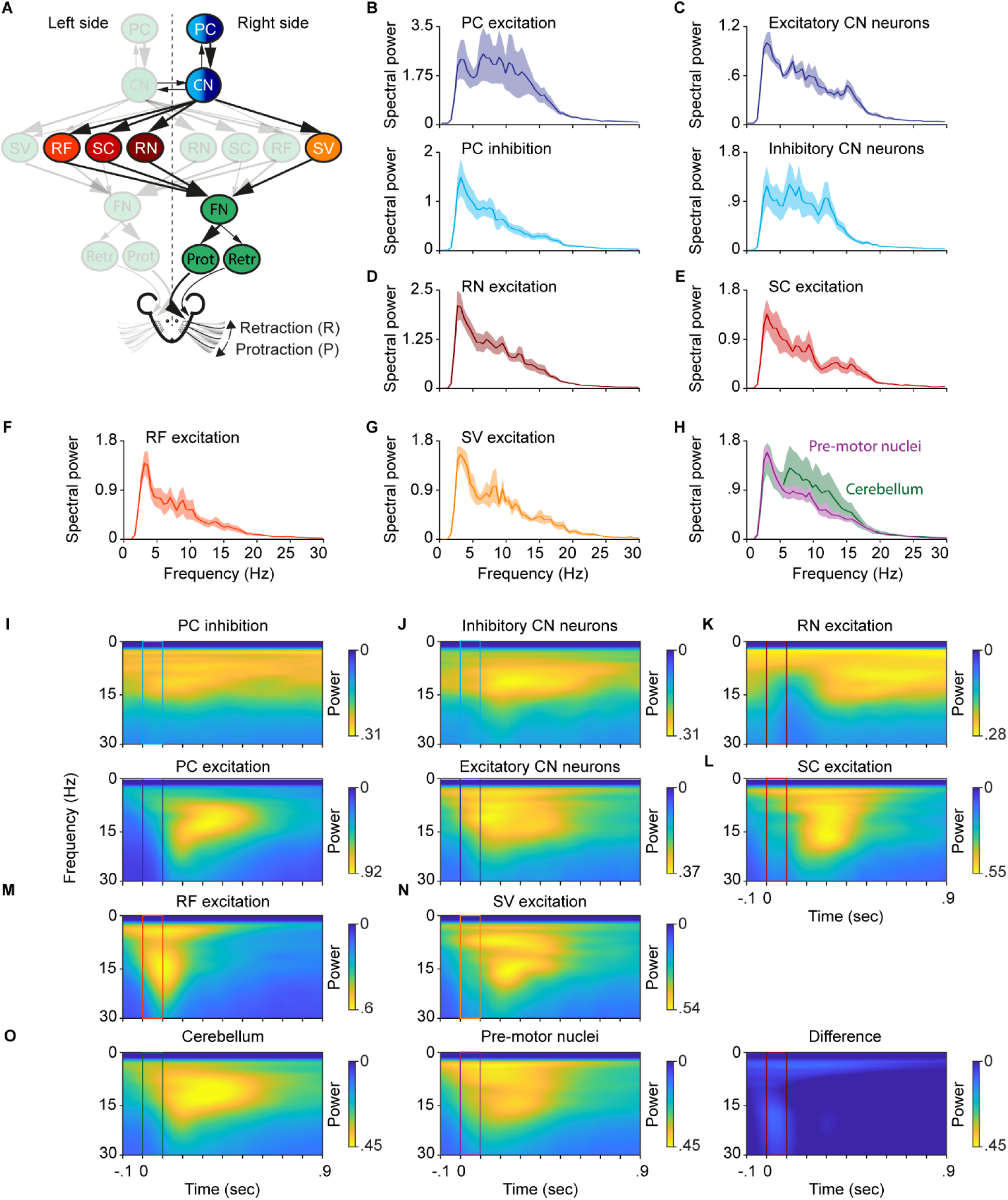
Cerebellar stimulation triggers whisker movements at a wide range of frequencies, resembling the combined effects of all its downstream targets. (**A**) A schematic of the neural circuit with colours indicating the sites of stimulation. (**B**) Top: Optogenetic inhibition of PCs induced whisker movements preferred in two frequency ranges: around 5 Hz and 20 Hz. Bottom: Optogenetic excitation of PCs induced whisker movements in a broad frequency range: from 2 Hz to 20 Hz. (**C**) Similar to (**B**), it’s inhibition (top) and excitation (bottom) of CN neurons. (**D, E, F, G**) are the spectral power of induced whisker movements when optogenetic excitation of SV, RF, SC, and RN respectively. (**H**) The sum of the spectral power of induced whisker movements from optogenetic excitation of SV, RF, SC, and RN overlaid with the spectrum of whisker movement during cerebellar stimulation. (**I**) Top: average spectrogram of the right whisker movements around stimulation during PC inhibition, ranging from -0.1 to 0.9 seconds. Spectrogram is calculated using the continuous wavelet transform, and high and low powers are denoted by yellow and blue colors, respectively. Bottom: similar as top, but for PC excitation. (**J**), (**K**),(**L**),(**M**),(**N**), are similar as in (**I**), but for RN, SC, RF, and SV excitation, respectively. (**O**) Left: similar to (**I**), but the average of all cerebellar stimulations. Middle: same as left, but for all pre-motor nuclei stimulation sites. Right: Difference between cerebellar and pre-motor nuclei stimulation sites. Abbreviation: Purkinje cell of the Paramedian lobule (PC), cerebellar nuclei (CN), facial nucleus (FN), reticular formation (RF), superior colliculus (SC), spinal trigeminal nucleus (SV), the red nucleus (RN), protractor muscles (Prot), retractor muscle (Retr), angle whisker position (Pos).

The statistical significance between values of the mean amplitude, distance, and latency (Figures 1-6) was assessed by performing a paired Wilcoxon-signed rank test. Significant results are denoted by an *, **, or *** for p<0.05, p<0.01, and p<0.001, respectively. Not applicable is denoted by N/a when there was no or too little data for one of the two sides present. Bar plots represent the median and interquartile ranges, as well as individual datapoints.

The amplitude was determined as follows; we extracted the most extreme peak (maximum or minimum) within 500 ms after the onset of stimulation. Then the two adjacent peaks were taken as the start and end points of the window of interest. The maximum value minus the minimum value within this window of interest was taken as the amplitude. The distance was calculated as the sum of the absolute value of the differential of the whisker trace and represents the cumulative sum of how many degrees the whiskers travelled after the stimulus onset. The latency was calculated as the time difference between the onset of stimulation and the timepoint when the whisker angle was above or below the baseline plus or minus 3 times the standard deviation, respectively. The window of interest for the baseline and standard deviation was the period 1000 ms before the onset of stimulation. During this window of interest, the baseline was determined as the mean of the whisker trace.

During the calculations of the amplitude, distance, and latency, the mean whisker trace of every mouse was used, resulting in a single data point per mouse. For visualisation, whisker traces were either high-pass filtered >2 Hz or normalised to the mean of 750 ms to 250 ms before stimulus onset.

### 2.8 Spectral analysis

Spectral analysis (Figure 7) was performed using custom-written MATLAB scripts. To calculate the power spectrum density estimate (PSD) shown in Figure 7 B-H, we created segments around stimulations from stimulation onset to 2 seconds after stimulation onset (preventing overlap between segments) and calculated the PSD using MATLAB’s periodogram function. Each segment was windowed with a Hamming window matching the segment length. A 2048 point discrete Fourier transform was used (nfft = 2048). Each segment resulted in a PSD, which we used to calculate the average PSD for individual animals. Afterwards, these averages were used to visualise the mean ± SEM for Figures 7 B-H. In addition, we created spectrograms from the whisker vector by calculating the continuous wavelet transform (CWT) using MATLAB’s function cwt using the default settings unless noted otherwise. The CWT was calculated from 0 to 30 Hz, as this is the range of frequencies in which mice spontaneously whisk (Bauer et al., 2022). The CWT gives information about the power of frequencies in the time-frequency domain and as the frequency vector is on a logarithmic scale, we interpolated this from 0 to 30 with a stepsize of 0.1 (Hz). Figures 7 I-L shows the average spectrogram around the stimulation. To create these spectrograms, we again created segments around the stimulation from -0.1 to 0.9 seconds and calculated the average spectrogram for every individual mouse, which were then used to calculate the average spectrogram per location of stimulation as visualized in 7 I-L.

## 3. Results

### 3.1 The activation of Purkinje cells, rather than their inhibition, triggers a highly rhythmic pattern of bilateral whisker movement

To compare whisker movements induced by optogenetic excitation versus inhibition of PCs, we applied 100 ms optogenetic stimulation to PCs in the medial part of the right paramedian lobule (PML) (Bauer et al., 2022; Zhai et al., 2024) while recording bilateral whisker movements of awake head-fixed mice. To up-regulate or down-regulate PC activity, we used genetic mouse lines that selectively express channelrhodopsin-2 (mouse-line Ai27D, n=7) and halorhodopsin (mouse-line Ai39, n=6) in PCs, respectively (Bauer et al., 2022; Han, Chen, Khan, Guo, & Regehr, 2020; Proville et al., 2014; Tsubota, Ohashi, Tamura, Sato, & Miyashita, 2011; Witter, Canto, Hoogland, de Gruijl, & De Zeeuw, 2013). Optogenetic excitation results consistently in rhythmic bilateral whisking (Figure 1C) with multiple cycles of whisking (Figures 1C and E), which is consistent over 7 mice (Figures 1G). In contrast, inhibiting PCs results in non-consistent rhythmic whisking on the right side with a smaller amplitude compared to excitation. Furthermore, inhibition of PCs does not result in whisking on the left side (Figures 1D, F, and H). To compare the effect of optogenetic stimulation on the right versus the left side, we quantified the maximal amplitude, the total distance covered during the induced movement, and the latency between the onset of stimulation and the onset of movement. During optogenetic excitation, no significant differences were found in amplitude, distance, and latency (Left: amplitude 9.56° (IQR 6.72–13.13°); distance 67.82° (IQR 62.38–75.20°); latency 40 ms (IQR 19.5–60.8 ms). Right: amplitude 13.99° (IQR 12.39–16.62°); distance 60.94° (IQR 57.16–90.03°); latency 25 ms (IQR 18.3–32.5 ms), median (25th–75th percentiles) Figure 1I). During optogenetic inhibition of PCs, the amplitude, distance, and onset likewise showed no significant differences between the left and right sides (Left: amplitude 1.05° (IQR 0.89–2.03°); distance 14.81° (IQR 11.41– 17.73°); latency 173 ms (IQR 153.5–215 ms). Right: amplitude 1.63° (IQR 1.20–1.79°); distance 18.45° (IQR 15.49– 19.66°); latency 38 ms (IQR 28.3–67.3 ms), median (25th–75th percentiles) Figure 1J). Together, these results show that only the upregulation of simple spike activity due to PC excitation elicits vigorous bilateral symmetric whisker motor responses. These whisker responses consist of a stereotyped rhythmic pattern which partly resembles the patterns observed during natural rhythmic whisking (Bauer et al., 2022; Zhai et al., 2024).

### 3.2 Neither stimulation of excitatory nor inhibitory cerebellar nuclear neurons alone consistently triggers a rhythmic pattern of bilateral whisker movement

PCs are the sole output of the cerebellar cortex and inhibit the CN. To investigate the role of excitatory and inhibitory CN on whisker movements, we selectively stimulated excitatory or inhibitory CN on the right side after injecting AAV9-Syn-FLEX-ChrimsonR-tdTomato into 7 VGluT2-ires-Cre or 8 Gad2-ires-Cre mice, respectively (Wang, Yu, Ren, De Zeeuw, & Gao, 2020) (Figure 2A and B).

Optogenetic stimulation of excitatory CN evokes an initial asymmetric bilateral movement (Figure 2C). The right and left whiskers protract and retract, respectively, followed by a bilateral protraction around the offset of the stimulus. Despite some inter-trial variability, the behaviour was relatively consistent over multiple trials (Figure 2D and F) and visible at the population level. The second protraction on the right side shows quite some variability (Figure 2H). Optogenetically stimulating inhibitory CN results in non-consistent bilateral movement (Figures 2C and 2E), mainly occurring around the stimulation offset (Figures 2E and G). At the population level, there was extensive variability (Figure 2I), suggesting that inhibiting CN can induce movement, but not in a stereotypical manner.

We compared the effect of stimulation on the whisking amplitude, distance covered by the whisker, and latency to the onset of movement. The whisking amplitude, whisking distance, and latency after stimulation of excitatory CN showed no significant difference between the left and right sides (Left: amplitude 3.13° (IQR 1.38– 4.09°); distance 24.77° (IQR 18.94–40.21°); latency 44 ms (IQR 36–71 ms). Right: amplitude 2.73° (IQR 2.21–4.54°); distance 31.58° (IQR 23.27–36.23°); latency 63.5 ms (IQR 55–76 ms), median (25th–75th percentiles)). Stimulation of the inhibitory CN likewise showed no significant difference between the amplitude and the distance covered by the whiskers between the left and right side (Left: amplitude 1.68° (IQR 1.11–2.24°); distance 18.77° (IQR 13.77–21.59°); latency 224 ms (IQR 157–359 ms). Right: amplitude 1.24° (IQR 1.14–1.57°); distance 16.82° (IQR 14.65–21.41°); latency N/a, median (25th–75th percentiles)). The latency could not be calculated for the right side after stimulation of the inhibitory CN. Overall, activation of excitatory CN can trigger non-stereotypical whisker movements, whereas activation of inhibitory CN does not trigger a rhythmic pattern of bilateral whisker movements.

### 3.3 Neuronal excitation in the red nucleus triggers bilaterally symmetrical non-rhythmic whisker movement

The RN is a pre-motor nucleus receiving projections from the contralateral CN (Figure 3A)(Novello et al., 2022) and its projections to the FN again cross the midline. To investigate the pathway downstream of the right cerebellar cortex, we stimulated the left RN. Previous electrical stimulations of the RN unreliably induced whisker movement in anaesthetized rats (Isokawa-Akesson & Komisaruk, 1987). To investigate how the RN affects whisker movement in the awake animal, we first injected AAV-ChrimsonR virus into the left RN (Figure 3B) in 9 mice which allows for modulation of the RN activity. Optogenetic stimulation in the RN results consistently in a prolonged protraction on the right side (Figures 3C, D, and E) which was also visible at the population level (Figure 3F). The whiskers on the left side also protracted, although with more variation in the population (Figure 3E). In addition, both the amplitude and distance covered by the whiskers did not show a significant difference (Left: amplitude 2.41° (IQR 2.01–4.48°); distance 18.90° (IQR 17.03–23.17°); latency 35 ms (IQR 5.5–59.8 ms). Right: amplitude 2.26° (IQR 1.67–2.97°); distance 18.00° (IQR 16.43–20.41°); latency 36.5 ms (IQR 16–241.5 ms), median (25th–75th percentiles) Figure 3G). The latency, although having a big difference between the mean, likewise showed no significant difference (left: latency 37.9 ± 12.7 ms, right: latency 130.6 ± 56.5 ms, mean ± SEM). Together, these results show that stimulation of the RN results in bilateral symmetric non-rhythmic whisking with a prolonged change in setpoint, suggesting that the RN could induce the symmetric changes of the midpoint during whisking behavior.

### 3.4 Neuronal excitation in the superior colliculus triggers bilaterally symmetrical quasi-rhythmic whisker movement

The superior colliculus (SC) projects directly to the facial nucleus (Figure 4A)(Bosman et al., 2011) and is considered to be a key area controlling mobile sensory organs such as the eyes, ears, and whiskers (May, 2006). 2-second electrical stimulation of the superior colliculus (50-1000 Hz) induces prolonged protraction (Hemelt & Keller, 2008) and a lesion of the SC results in a more retracted resting position of the contralateral whisker (Kaneshige, Shibata, Matsubayashi, Mitani, & Furuta, 2018). To optogenetically investigate the role of the SC on whisking, we injected AAV-ChrimsonR virus into the left SC of 7 mice (Figure 4B) and optogenetically stimulated the SC for 100 ms.

Optogenetic stimulation of the SC resulted in an initial protraction on the right side (Figure 4C), which is consistent over multiple trials (Figure 4D) and was evident in the average whisker trace around stimulation of multiple trials (Figure 4E). On a population level, there was quite some variability regarding the response on both the left and right sides, except for the prolonged protraction (Figure 4F). Overall, at a population level, we induced a prolonged bilateral symmetrical movement with superimposed faster whisking cycles. Comparing the amplitude, distance covered by whiskers and latency between the left and right sides showed that there is no significant difference (Left: amplitude 4.38° (IQR 2.89–5.55°); distance 29.81° (IQR 18.81–40.72°); latency 88 ms (IQR 45.5–174.5 ms). Right: amplitude 4.19° (IQR 3.29–6.23°); distance 39.60° (IQR 28.66–46.79°); latency 21 ms (IQR 15–35 ms), median (25th–75th percentiles), Figure 4G). These results suggest that the SC could regulate bilateral and symmetrical changes of the midpoint of the whiskers in a quasi-rhythmic fashion with a lasting effect after stimulation offset.

### 3.5 Neuronal excitation in the reticular formation triggers a fast bilateral whisker movement

The RF and more specifically, the intermediate part of the reticular formation, has received the most attention in previous studies focusing on the neural circuit behind whisking (Deschenes, Kurnikova, Elbaz, & Kleinfeld, 2016; Golomb et al., 2022; Kleinfeld et al., 2014; Moore et al., 2013; Takatoh et al., 2022). Electrolytic lesioning of the RF abolishes whisking and kainic acid injection in anaesthetized rats around the RF can induce whisking around 10 Hz (Moore et al., 2013). The afferents of the FN from the RF have been described as bilateral (Figure 5B) (Hattox et al., 2002). To investigate the role of the RF in awake animals, we injected AAV-ChrimsonR virus in the left RF (Figure 5B) and applied 100 ms optogenetic stimulation to the RF. Soon after the onset of optogenetic stimulation, the mouse protracted the right-side whiskers and retracted the left side (Figure 5C). These initial asymmetrical movements were followed by other fast whisking cycles on the right side. These whisker patterns were consistent over multiple trials (Figure 5D-E) and for 4 mice (Figure 5F). The amplitude of the right side was almost twice the amplitude of the left side (left: 4.88 (IQR 3.72–6.93°) ; right: 13.16° (IQR 7.47–14.85°), median (25th–75th percentiles), Figure 5G); however, this difference was not significant (p>0.05). The distance covered by the whiskers likewise did not show a significant difference, while their median were 67.82° (IQR 62.38–75.20°) and 60.94° (IQR 57.16–90.03°) for the left and right sides, respectively. The short latency for the initial bilateral asymmetrical response was 40 ms (IQR 19.5–60.8 ms) and 25 ms (IQR 18.3–32.5 ms) for the left and right sides, respectively. Together, these results confirm the involvement of the RF in generating the fast-whisking component. In addition, this short latency bilateral asymmetrical whisker response highlights the immediate effect of the RF on the FN and the whiskers.

### 3.6 Neuronal excitation in the spinal trigeminal triggers slow bilateral whisker movement

The SV receives projections from the cerebellum (Figure 6A) and is also part of a reflex arc, receiving direct projections from the ipsilateral trigeminal nerve (Matthews et al., 2015). To investigate its contribution to whisking behaviour, we injected an AAV-ChrimsonR virus in the right SV (Figure 6B) of 7 mice, enabling modulation of its activity. Upon SV stimulation, we observed a response of the whiskers in opposite directions, the left and right sides initially retract and protract, respectively (Figures 6C, D, and E). The right side showed 2 whisking cycles (Figure 6C). These first and second responses were consistent over all trials within one mouse (Figure 6D). At the population level, the first response was consistent, while the second response was more variable, although most animals showed a prolonged bilateral protraction (Figure 6E). Although in opposite directions, the amplitude of the left and right sides was similar (5.23° (IQR 3.60–9.85°) and 6.00° (IQR 2.95– 9.66°), respectively, median (25th–75th percentiles), Figure 6G). The movement of the whiskers on the left and right side consisted of one and two cycles, respectively; however, this did not result in a difference in distance covered by the whiskers (47.56° (IQR 29.87–67.43°) and 54.40° (IQR 24.62–63.65°), respectively, median (25th– 75th percentiles), Figure 6G). Furthermore, the latencies between the left and right sides were different, although not significant (58.5 ms (IQR 45–109 ms) and 17 ms (IQR 8–43 ms), respectively, median (25th–75th percentiles), Figure 6G). Together, these results show that excitation of the SV results in an asymmetrical initial response, followed by a prolonged bilateral protraction.

### 3.7 Cerebellar stimulation triggers whisker movements at a wide range of frequencies, resembling the combined effects of all its downstream targets

In our experimental condition, voluntary whisking behaviour occurs mostly within a range of 2-25 Hz (Bauer et al., 2022). To test at which frequencies mice whisk during optogenetic stimulation of brain areas involved in this study (Figure 7A), we computed the power spectra of whisking around stimulation of these areas. We focused on the right whiskers as whisker kinematics did not differ between the left and right sites for amplitude, distance covered, and onset. In addition, whisker movements can be split into the ‘slow component’, more related to the midpoint (<6 Hz) and its faster component (>6 Hz), associated with the phase of whisking (Chen, Augustine, & Chadderton, 2016; Zhai et al., 2024). To include both, we filtered the whisker trace between 2-50 Hz. Inhibition and excitation of PCs resulted in whisking at frequencies ranging from 2-20 Hz (Figure 7B), covering almost the complete range of frequencies observed during voluntary whisking (Bauer et al., 2022). In addition, the power of the frequencies observed during excitation of PCs is substantially higher compared to the power of the frequencies observed during inhibition of PCs, in line with the results in Figure 1.

Stimulation of CN resulted likewise in whisking behaviour within the range of 2-20 Hz (Figure 7C), however, stimulation of inhibitory CN shows peaks at 3, 6, and 12 Hz, while stimulation of excitatory CN shows its peaks at 3, 7, and 15 Hz.

Stimulation of the RN evoked whisker movement mainly at low frequencies (Figure 7D) with a single peak occurring around 2 Hz, highlighting the prolonged protraction. Stimulation of the SC results in peaks at 9 and 15 Hz (Figure 7E). The RF has been revealed as a key area in the oscillator circuit for whisking (Moore et al., 2013; Takatoh et al., 2022). During our 100 ms stimulation of the RF, we observed whisking at frequencies ranging from 3-20 Hz, with its main peaks at 2 and 9 Hz. Similar to what we observed for the SC, the power spectrum of whisking behaviour during stimulation of the SV showed peaks at 8 and 15 Hz (Figure 7G).

Since the output of PCs passes via the CN through the RN, SC, RF, and SV before it reaches the FN, we tested whether it could be a pattern separation downstream of the cerebellum in terms of frequencies. Summing up the power spectrum of whisking behaviour during stimulation of the RN, SC, RF, and SV resulted in a power spectrum covering frequencies from 3-25 Hz, greatly overlapping the whisking frequencies observed during the stimulations in the cerebellum (PC excitation, PC inhibition, stimulation of excitatory, and inhibitory CN) (Figure 7H). These results show that the RN, SC, RF, and SV induce whisking at different frequencies, which cover the range of frequencies observed during both natural whisking and the combination of stimulations we performed on the cerebellum.

The Fourier transform allows for investigation solely in the frequency domain. To additionally investigate the time-frequency domain, we applied a continuous wavelet transform to the same whisker traces aligned to stimulation onset. This method allows for the examination of frequency-specific power over time. PC inhibition resulted in a band of elevated power predominantly in the low frequency ranges, whereas PC excitation produced high power at higher frequencies after stimulation offset, persisting for around 500 ms (Figure 7I). Stimulation of inhibitory CN neurons elicited a power increase at 7-15 Hz from stimulation onset and lasted around 700 ms. The effect of stimulating excitatory CN neurons resulted in a high power in a broader frequency range from 2-16 Hz, again from stimulation onset and lasting for about 500 ms (Figure 7J). The spectrogram of RN stimulation showed that apart from its low frequency component (∼3-5 HZ), the higher frequencies emerged only later, 200 ms after stimulation onset, but lasted for more than 700 ms (Figure 7K). SC excitation initially resulted in low frequency power (∼3 HZ), and 200 ms after stimulation onset, enhanced power in higher frequencies up to 20 Hz (Figure 7L). RF excitation produced a distinct and transient increase in power across 2– 20 Hz for approximately 200 ms from stimulation onset (Figure 7M). In contrast, SV excitation showed two components: a lower frequency component of 5-10 Hz emerging at stimulation onset, and a second component of 11-20 Hz, which initiated around stimulation offset. Both components lasted up to 500 ms after stimulation onset (Figure 7N). To compare the effects of cerebellar versus pre-motor nuclear stimulation, we computed average spectrograms aligned to stimulation onset for PC excitation, PC inhibition, and excitatory and inhibitory CN stimulation. These conditions resulted in increased power across nearly the entire natural whisking frequency range, starting at stimulation onset and extending up to 600 ms. Similarly, averaging spectrograms from RN, SC, RF, and SV excitation revealed a broad increase in power across frequencies beginning at stimulation onset and lasting up to 600 ms. To directly contrast the two groups, we subtracted the averaged cerebellar spectrograms from the pre-motor spectrograms. This subtraction largely abolished the power increase surrounding the stimulation period (Figure 7O), suggesting that, beyond frequency domain differences, a temporal pattern separation is also evident in the time-frequency domain between cerebellar and downstream pre-motor nuclei. We propose that the cerebellum targets all these pre-motor nuclei to fine-tune the various patterns of whisking behaviour.

### 3.8 Whisker movements are predominantly symmetrical with a positive cross-correlation

To capture more dynamics of the movements, we calculated the cross-correlation between the left and right whiskers during the stimulation (0 – 100 ms) and a post-stimulus (100 – 400 ms) period. From this cross-correlation, we extracted the maximal cross-correlation and its corresponding lag. For all stimulation sites the peak of most common lag was at lag 0. In addition, although perhaps unexpected due to initial (small) opposite movements for stimulation of the RF and SV (Figures 5 and 6, respectively), all stimulation sites showed a predominantly positive cross-correlation, highlighting that most of the triggered whisker movements in these periods were bilateral and symmetrical (Figure S1).

## 4. Discussion

In this study, we investigated the whisker movement kinematics resulting from the stimulation of several neuronal types in the cerebellum and in the downstream brainstem nuclei connecting the cerebellum to the FN. We found that different pre-motor nuclei induce distinct patterns of whisking behaviour, as they contribute to different components of movement kinematics, suggesting that pattern separation occurs downstream of the cerebellum. Optogenetic excitation of PCs in the medial PML resulted in long-lasting rhythmic whisker movements on both sides (Figure 1), consistent with our recent findings for the ipsilateral side (Zhai et al., 2024). This pattern of whisker movements is quite different from the whisker movement during the excitation of PCs from other cerebellar lobules (Romano et al., 2020; Zhai et al., 2024) or during shorter stimulations (Bauer et al., 2022). The power spectrum of these whisker movements consisted of multiple peaks at different frequencies, ranging from ∼2-20 Hz. This range includes both whisker midpoint and fast rhythmic components, which have been previously separated using a cut-off frequency of 6 Hz filter (Chen et al., 2016; Cheung, Maire, Kim, Sy, & Hires, 2019). A similar range of whisking could be evoked, inhibiting the same medial PML area, although this evoked smaller and unilateral whisker movements compared to excitation. This smaller ipsilateral movement, however, was initiated after a shorter time delay relative to the light stimulus onset. This is similar to what has been shown for forelimb movement (Lee et al., 2015) and is consistent with the “disinhibition hypothesis,” stating that transiently reducing the firing rate of PCs can activate motor areas via disinhibition of the CN (Ito, 1984, 2001; Witter et al., 2013). Such a mechanism is also in line with the disinhibition of CN neurons, correlating well with eyelid kinematics (Heiney, Kim, Augustine, & Medina, 2014; Ten Brinke, Boele, & De Zeeuw, 2019). Stimulation of excitatory and inhibitory CN evoked similar patterns of bilateral whisker movements (Figure 2), which were different from those induced by PCs stimulation (Figure 1). Looking at the latency of whisker movement, stimulation of inhibitory CN neurons can evoke an earlier onset of contralateral whisking compared to excitation, while stimulation of inhibitory CN evokes a shorter latency of ipsilateral (relative to CN) whisking, possibly due to different downstream pathways. Overall, CN stimulation did not yield the complex whisking pattern induced by PC excitation. One possibility is that PC excitation affects both excitatory and inhibitory CN neurons, and the coordinated activation of both is required to generate a clear pattern of rhythmic whisking.

The CN project to numerous areas of the central nervous system (T. M. Teune, J. van der Burg, J. van der Moer, J. Voogd, & T. J. Ruigrok, 2000), among which are those strongly connected to the motoneurons that control whisking activity in the FN and receive projections from the whisker motor cortex (Hattox et al., 2002). One brain area for which the direct anatomical connection to the FN is still controversial is the RN. For this brain area, it was also unclear whether its stimulation could induce whisker movement (Isokawa-Akesson & Komisaruk, 1987). Here, using optogenetic stimulation, we show for the first time that excitation of the neurons in the RN of awake mice reliably induces whisker movements, which are more prominent on the contralateral (relative to the RN, Figure 3). Whether this is the result of a monosynaptic connection to the FN (Hattox et al., 2002) or a disynaptic connection via the RF (Takatoh et al., 2013) remains unclear. It is possible that the output generated by our stimulation reached the FN via multiple interconnected pathways. Despite that, the whisker movement induced by the RN stimulation consisted of a single, relatively slow whisking cycle, which was remarkably unique.

Optogenetic stimulation of the SC, instead, resulted in a bilateral protracted baseline position with superimposed a faster component (Figure 4). These results are in line with previous studies showing that electrical stimulation of the SC results in a more protracted contralateral whisker (Hemelt & Keller, 2008) and that lesioning the SC results in a more retracted contralateral whisker (Kaneshige et al., 2018).

Previous studies in lightly anaesthetized rats showed that the activity of a group of neurons in the intermediate portion of the RF is phase-locked to protraction and that stimulation with the kainic acid injection of this area results in rhythmic whisking (Kleinfeld et al., 2014; Moore et al., 2013). These studies were followed by another that clarified the mechanism of rhythmogenesis in this brain area, which is thought to be the central pattern generator of oscillatory whisking (Takatoh et al., 2022). Our stimulation of the RF, lasting 100 ms, resulted in a single initial asymmetric whisk cycle followed by a big protraction on the contralateral side (relative to the RF) (Figure 5). Such asymmetrical bilateral response, in natural circumstances, can be predictive of head and body movements, such as turning during walking (Sofroniew et al., 2014). Therefore, despite the RF being described as the central pattern generator (Moore et al., 2013; Takatoh et al., 2022), our 100 ms optogenetic stimulation did not induce multiple whisking cycles as PC, CN and SC excitation.

Finally, we investigated the SV, which is involved in a feedback loop receiving sensory input directly from the whisker follicle and directly innervating the FN to modulate motor neurons (Matthews et al., 2015). Stimulation of this area also resulted in asymmetric whisker movements with a short latency for the ipsilateral side (Figure 6). Interestingly, the movements consisted of two cycles on the ipsilateral (relative to the SV) and only one slower cycle on the contralateral (relative to the SV) side.

We compared the frequencies at which mice whisked during cerebellar optogenetic stimulation and stimulation of pre-motor nuclei downstream of the cerebellum. During both types of stimulation, we found that they together comprise the whisking frequencies seen during spontaneous whisking (2-20 Hz). This overlap in frequencies was not only present in the frequency domain, but also in the time-frequency domain as subtracting the average spectrogram of cerebellar stimulation from the pre-motor nuclei stimulation resulted in a spectrogram where all frequency components were abolished.

In addition, we investigated more dynamics of movements by calculating the cross-correlation between the movements of the left and right whiskers. This resulted in predominantly positive cross-correlations for all stimulation sites, which is in line with our previous finding during spontaneous whisking, where the most probable phase difference is 0 (Romano et al., 2022).

Taken together, our results suggest these pre-motor whisker nuclei contribute to distinct aspects of whisking kinematics. We propose that there is a pattern separation from the efferent of the cerebellar cortex in the downstream pre-motor whisker nuclei. Because in our experiments we used head-fixed mice, we were unable to determine if some pre-motor nuclei induced predictive whisker movements preceding head and body movements.

In conclusion, our systematic study on the brain-wide network downstream of the cerebellar cortex that controls whisker movements shows, for the first time, the complexity of the whisking patterns induced by manipulating the main hubs of the network controlling whiskers in rodents. The neuronal activity of the upstream PC, therefore might tune precise whisker movement by differently modulating the activity of its downstream target in a highly coordinated manner.

## Acknowledgements

We thank E.D. Haasdijk (Department of Neuroscience, Erasmus MC) for technical assistance.

## Author Contributions

S.B. and P.Z. contributed equally to this work.

S.B. P.Z. N.v.W. V.R.

Conceptualization: P.Z. and V.R. Data curation: S.B., P.Z., and V.R. Formal analysis: S.B., P.Z., and V.R. Funding acquisition: V.R. (Veni) S.B. (DBI2) Investigation: P.Z., N.v.W., and V.R. Methodology: S.B., P.Z., N.v.W., and V.R. Project administration: N.v.W. and V.R. Resources: N.v.W. Software: S.B. and P.Z. Supervision: V.R. Visualization: S.B. and P.Z. Writing – original draft: S.B., P.Z., and V.R. Writing – review & editing: S.B., P.Z., and V.R.

## Data Accessibility Statement

All data are available from the Lead Contact upon request. The custom code complementing BWTT whisker tracking can be obtained via https://github.com/elifesciences-publications/BWTT_PP. The custom code WhiskEras can be obtained via (https://gitlab.com/c7859/neurocomputing-lab/whisker/whiskeras-2.0) and more information is available at (https://whiskeras.nl/).

## Abbreviations

PC: Purkinje cell of the Paramedian lobule
CN: cerebellar nuclei
FN: facial nucleus
RF: reticular formation
SC: superior colliculus
SV: spinal trigeminal nucleus
RN: the red nucleus
Prot: protractor muscles
Retr: retractor muscle
Pos: angle whisker position

---

**Figure S1:**
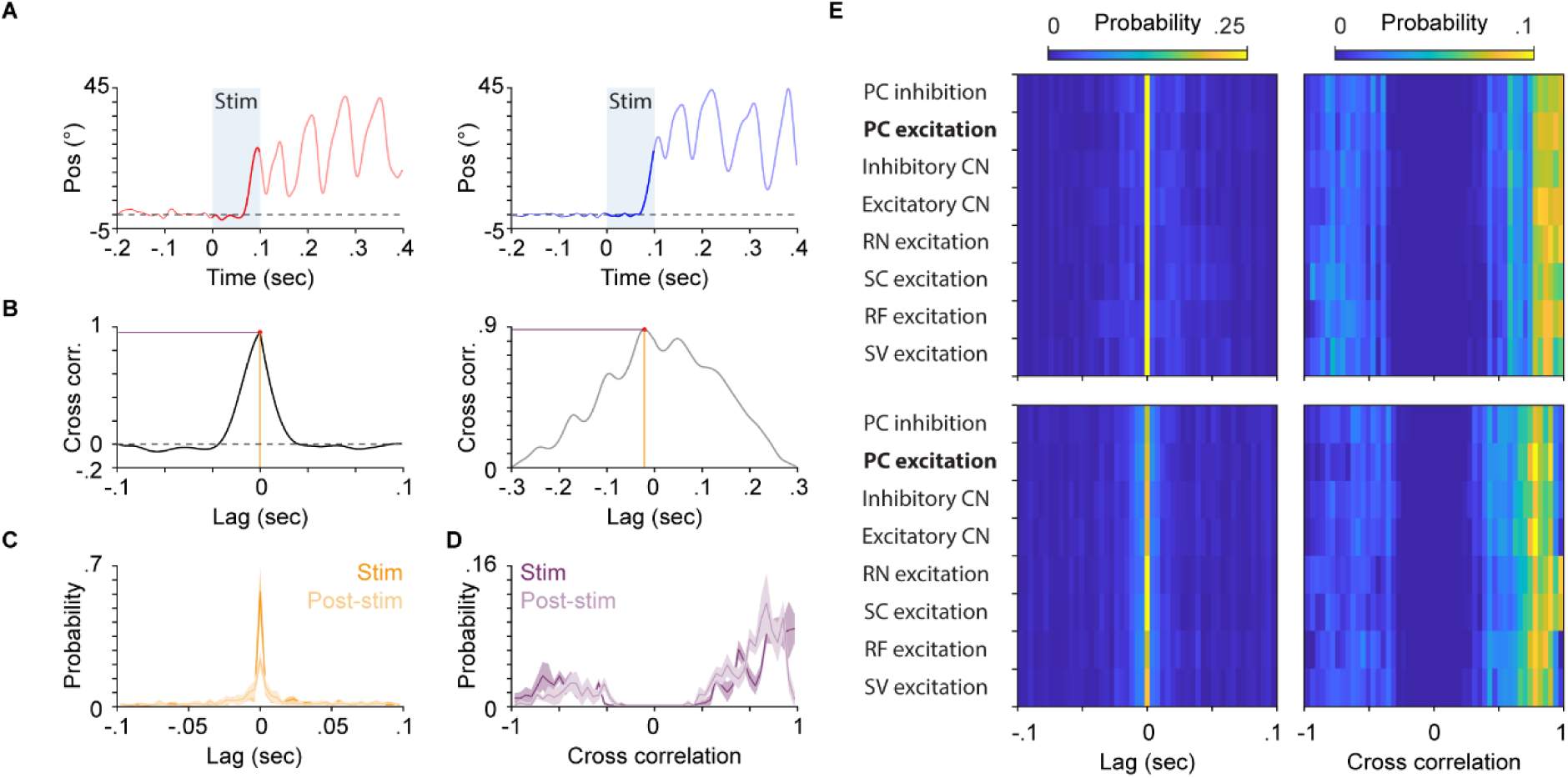
Left and right whisker movements are predominantly symmetrical with a positive cross-correlation. (**A**) Left: example whisker trace of left whisker during PC excitation. Stimulus lasts from 0-100 ms, which highlighted by the blue shaded area and the thick red line. The post-stimulus time is highlighted by the lighter colored thick red line from 100 – 400 ms after stimulus onset. Right: similar, but for the right whisker during the same PC excitation. (**B**) Left: cross-correlation between left and right whisker traces during the stimulation period from panel (**A**). The maximal cross-correlation is highlighted by a red dot and a purple horizontal line. The corresponding lag is highlighted by a red dot and a yellow vertical line. Right: Similar to the left side, but for the post-stimulus period of 100 – 400 ms after stimulus onset. (**C**) Probability distribution of all the lags corresponding to the maximal cross-correlation as indicated in (**C**) for one exemplary mouse. Dark and light yellow indicate during stimulus (0 – 100 ms) and post-stimulus (100 – 400 ms), respectively. (**D**) Probability distribution of all maximal cross-correlations as indicated in (**C**) for one exemplary mouse. Dark and light purple indicate during stimulus (0 – 100 ms) and post-stimulus (100 – 400 ms), respectively. (**E**) Top left: Average probability distribution of lags during stimulus, similar to (**C**) but for all stimulation sites. Blue and yellow indicate low and higher probabilities, respectively. Top Right: Average probability distribution of cross-correlations during stimulus, similar to (**D**) but for all stimulation sites. Blue and yellow indicate low and higher probabilities, respectively. Bottom: similar to the top part, but during the post-stimulus period.

**Figure S2:**
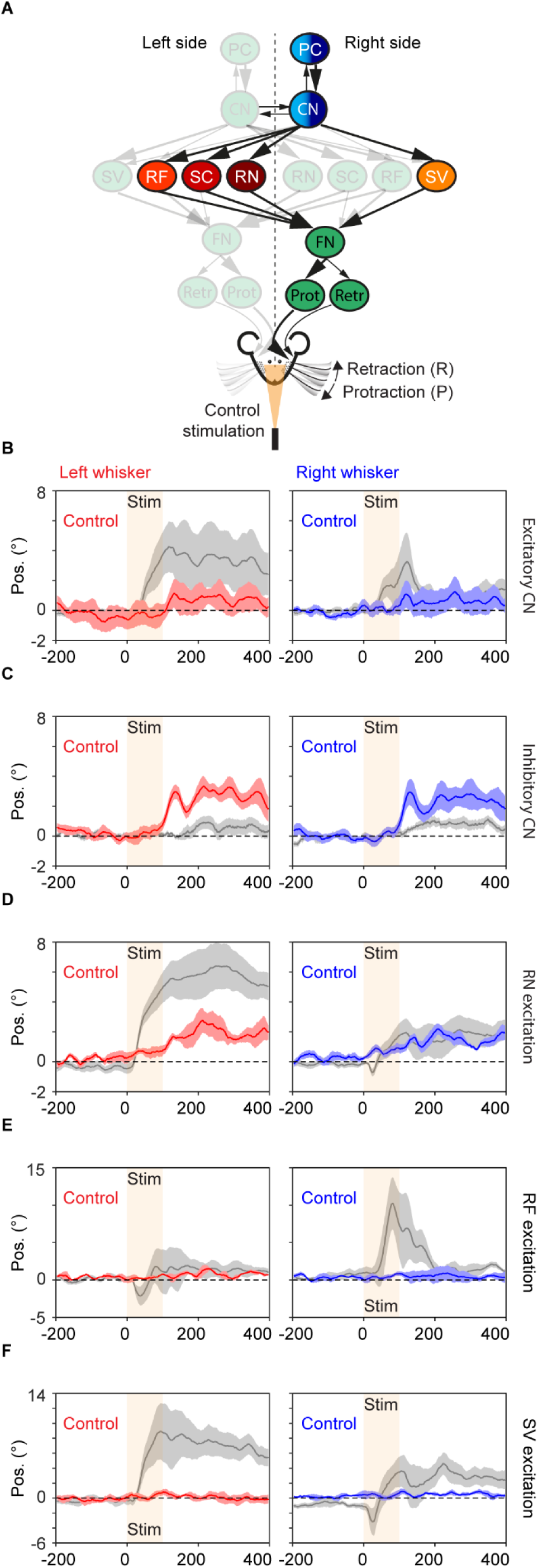
(**A**) A schematic of the neural circuit with colours indicating the sites of stimulation. The orange light given as a control (visual) stimulus was placed in front of the mouse as indicated in the schematic. (**B**) Left: Average whisker movement of the left whisker aligned to either the real (stimulation of excitatory CN) or control stimulus in grey and red, respectively. The period of stimulation is highlighted by the orange shaded area. Right: Average whisker movement of the right whisker aligned to either the real (stimulation of excitatory CN) or control stimulus in grey and blue, respectively. The period of stimulation is highlighted by the orange shaded area. (**C**), (**D**), (**E**), and (**F**) are all similar to (**B**) but for the stimulation of inhibitory CN, RN excitation, RF excitation, and SV excitation, respectively. Abbreviation: Purkinje cell of the Paramedian lobule (PC), cerebellar nuclei (CN), facial nucleus (FN), reticular formation (RF), superior colliculus (SC), spinal trigeminal nucleus (SV), the red nucleus (RN), protractor muscles (Prot), retractor muscle (Retr), angle whisker position (Pos).

